# Targeting against HIV/HCV Co-infection using Machine Learning-based multitarget-quantitative structure-activity relationships (mt-QSAR) Methods

**DOI:** 10.1101/605162

**Authors:** Yu Wei, Wei Li, Tengfei Du, Zhangyong Hong, Jianping Lin

## Abstract

Co-infection between HIV-1 and HCV is common today in certain populations. However, treatment of co-infection is full of challenges with special consideration for potential hepatic safety and drug-drug interactions. Multitarget inhibitors with less toxicity may provide a promising therapeutic strategy for HIV/HCV co-infection. However, identification of one molecule acting on multiple targets simultaneously by experimental evaluation is costly and time-consuming. *In silico* target prediction tools provide more opportunities for the development of multitarget inhibitors. In this study, by combining naive Bayesian (NB) and support vector machine (SVM) algorithms with two types of molecular fingerprints (MACCS and ECFP6), 60 classification models were constructed to predict the active compounds toward 11 HIV-1 targets and 4 HCV targets based on the multitarget-quantitative structure-activity relationships (mt-QSAR). 5-fold cross-validation and test set validation was performed to confirm the performance of 60 classification models. Our results show that 60 mt-QSAR models appeared to have high classification accuracy in terms of ROC-AUC values ranging from 0.83 to 1 with a mean value of 0.97 for HIV-1 models, and ROC-AUC values ranging from 0.84 to 1 with a mean value of 0.96 for HCV. Furthermore, the 60 models were applied to comprehensively predict the potential targets for additional 46 compounds including 27 approved HIV-1 drugs, 10 approved HCV drugs and 9 selected compounds known to be active on one or more targets of HIV-1 or those of HCV. Finally, 18 hits including 7 HIV-1 approved drugs, 4 HCV approved drugs and 7 compounds were predicted to be HIV/HCV co-infection multitarget inhibitors. The reported bioactivity data confirmed that 7 compounds actually interacted with HIV-1 and HCV targets simultaneously with diverse binding affinities. Of those remaining predicted hits and chemical-protein interaction pairs involving the potential ability to suppress HIV/HCV co-infection deserve further investigation by experiments. This investigation shows that the mt-QSAR method is available to predict chemical-protein interaction for discovering multitarget inhibitors and provide a unique perspective on HIV/HCV co-infection treatment.

## 1. Introduction

Human immunodeficiency virus type-1 (HIV-1) is the causative factor of acquired immunodeficiency syndrome (AIDS), a pandemic disease[1]. In addition, hepatitis C virus (HCV) infection causes acute and chronic liver disease, including cirrhosis and hepatocellular carcinoma[2]. Unfortunately, since HIV-1 and HCV have similar routes of transmission, the risk of HIV/HCV co-infection is very high[3]. According to the World Health organization (WHO), about 2.3 million persons of the estimated 36.7 million living with HIV had been infected with HCV globally in 2015. It is necessary for those people to be diagnosed and provided with effective and reasonable treatment for both HIV and hepatitis as a priority[4]. However, in the treatment of HIV/HCV co-infection, special considerations must be given to potential drug-drug interactions and toxicities. The standard treatment for HIV-1 is highly active antiretroviral therapy (HAART), which improves the life quality of AIDS patients by reducing viral load[5]. HAART utilizes a cocktail of drugs to obstruct multiple viral proteins in the viral life cycle, including reverse transcriptase, integrase, and protease[6]. Although it is effective, HAART needs to sequentially administer single-target drugs that may result in drug-drug interactions, poor treatment adherence and the emergence of drug resistance. The current therapy for HIV/HCV co-infection includes injecting pegalated α-interferon (PEG-IFN) regularly plus taking Ribavirain (RBV) daily in combination with HAART[7]. Similar to the HARRT treatment regimen, additive drug toxicity, high cost and long-term therapy cycle should be taken into account during the process of HIV/HCV co-infection treatment. In order to avoid these undesirable factors, the conception of multitarget therapy using a single drug capable of inhibiting simultaneously two or more viral targets was proposed to reduce the complexity of the treatment[5].

With the development of computer-aided drug design (CADD) technology, several *in silico* methods have been exploited to predict the chemical-protein interaction (CPI) pairs between drugs and target proteins[8,9]. These methods can be approximately classified into two categories: structure-based and ligand-based, such as pharmacophore modeling[10], similarity searching[11,12] and molecular docking[13,14]. Conventional quantitative structure-activity relationship (QSAR) generally takes a group of analogs against one target into account to elucidate the relationship between chemical structure and biological activity. In 2009, Vina et al. developed a multitarget-QSAR (mt-QSAR) classification model to predict the probability of drugs binding more than 60 different receptors based on molecular connectivity and receptor sequences[15]. Furthermore, Jacob et al. proposed chemogenomic method to predict CPI[16], and Wang et al. used chemogenomic method to identify new ligands for four targets and the results was validated by experiments[17]. As a new interdisciplinary field, chemogenomics tries to completely match the target and ligand space, and determine available ligands for all targets[9]. Cheng et al. compared the CPI prediction performance between mt-QSAR and computational chemogenomic methods, and the result showed that mt-QSAR had a better performance than chemogenomic, which had a high risk of overfitting and a high false positive rate in the external validation set[18].

In this study, mt-QSAR method was applied to predict the chemical-protein interactions (CPIs) for 11 key targets related to HIV-1 and 4 targets related to HCV. The workflow is depicted in Fig 1. Firstly, combining two machine learning algorithms (naive Bayesian[19,20]and support vector machine[21]) and two molecular fingerprints (MACCS[22] and ECFP6[23]), we developed multiple mt-QSAR models for identifying inhibitors against 11 HIV-1 related and 4 HCV related targets. Secondly, 5-fold cross-validation and test set validation was used to evaluate the performance of all models. Thirdly, to verify the application of this strategy, the developed mt-QSAR models were employed to predict CPI for 27 approved HIV-1 drugs and 10 approved HCV drugs well as 9 compounds known to be active on one or more targets of HIV-1 or HCV. The predicted results shown 18 hits to be potential HIV/HCV co-infection multitarget inhibitors, which were further confirmed by reported bioactivity resulting in that 7 compounds actually interacted with HIV-1 and HCV targets simultaneously with diverse binding affinities. Our study indicates that machine learning-based mt-QSAR approaches could be potentially applied in discovery of HIV/HCV co-inhibitors and multitarget drug discovery.

**Fig 1.**
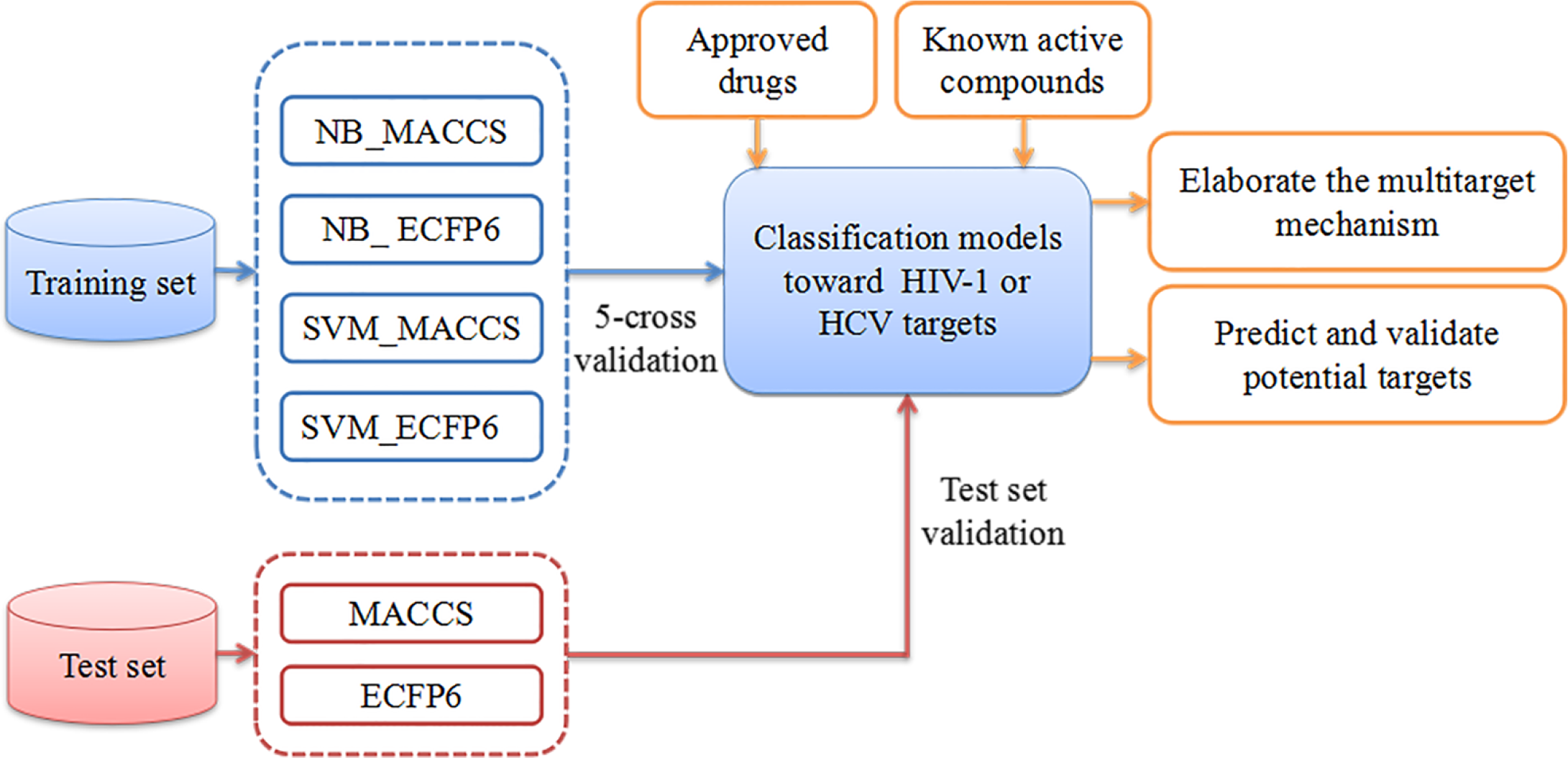
Flowchart of mt-QSAR models building process toward HIV-1/HCV co-infection (the blue, red and orange objects represent the phase of construction, validation and application of mt-QSAR models, respectively).

## 2. Results

### 2.1 Experimental datasets analysis

In this study, a total of 11 targets related to HIV-1 and 4 targets related to HCV were obtained. The number of known small molecule inhibitors that act on HIV-1 and HCV targets were 11,006 and 1,431, respectively. The number of active compounds for each target was shown in Fig 2. A total of 11 datasets were generated for HIV-1 system, including 157 inhibitors and 471 decoys for CXCR4, 370 inhibitors and 1,110 decoys for CCR5, 293 inhibitors and 879 decoys for GP120, 21 inhibitors and 63 decoys for GP41, 3,256 inhibitors and 9,768 decoys for RT, 2,215 inhibitors and 6,645 decoys for IN, 4,111 inhibitors and 12,333 decoys for PR, 101 inhibitors and 303 decoys for Gag-pol, 438 inhibitors and 1,314 decoys for protein Tat, 29 inhibitors and 87 decoys for PKC, 15 inhibitors and 45 decoys for CYP3A. There were 4 datasets for HCV system, including 470 inhibitors and 1,410 decoys for NS3/4A, 19 inhibitors and 57 decoys for NS4B, 40 inhibitors and 120 decoys for NS5A, 902 inhibitors and 2,706 decoys for NS5B. Each of 15 datasets above was divided into training set and test set.

**Fig 2.**
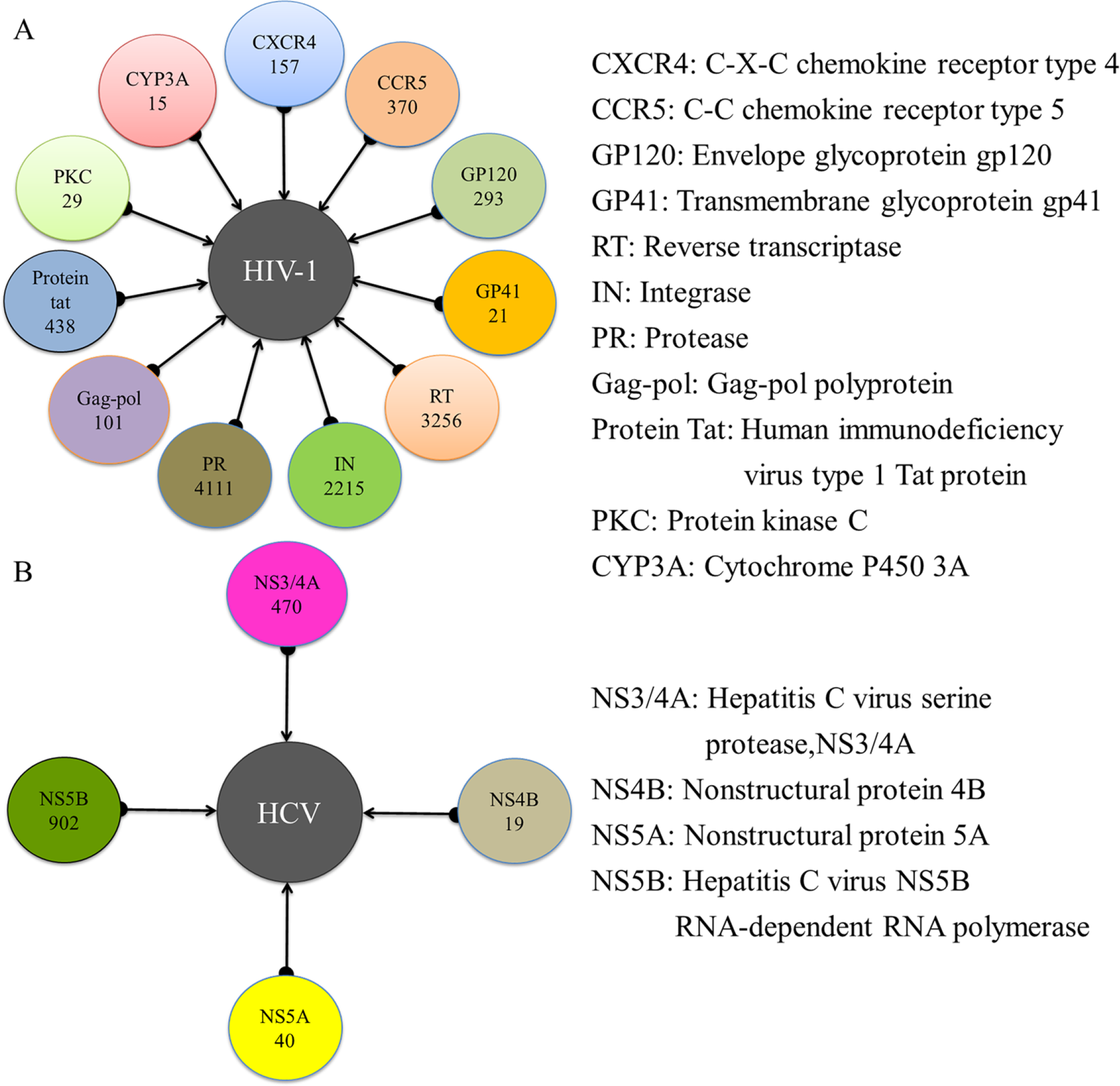
Summary of targets related to HIV-1 (A) and HCV (B) and the total number of active compounds for each target.

Before modeling implementation, Tanimoto similarity index, which is a generally-used measure for evaluating molecular similarity among numerous compounds, was used to evaluate fingerprint-based similarity using ECFP4 fingerprint[24,25]. The Tanimoto similarity index of a set of compounds in training set and test set for each of 15 datasets were computed, and the results were shown in Table 1. For HIV-1, the Tc value of training sets and test sets in 11 datasets range from 0.124 to 0.202 and 0.117 to 0.215, respectively. For HCV, the training sets and test sets of 4 datasets have the similar Tc values ranging from 0.141 to 0.182. Table 1 shows that compounds in the training sets and test sets of 15 datasets were sufficiently diverse.

**Table 1.**
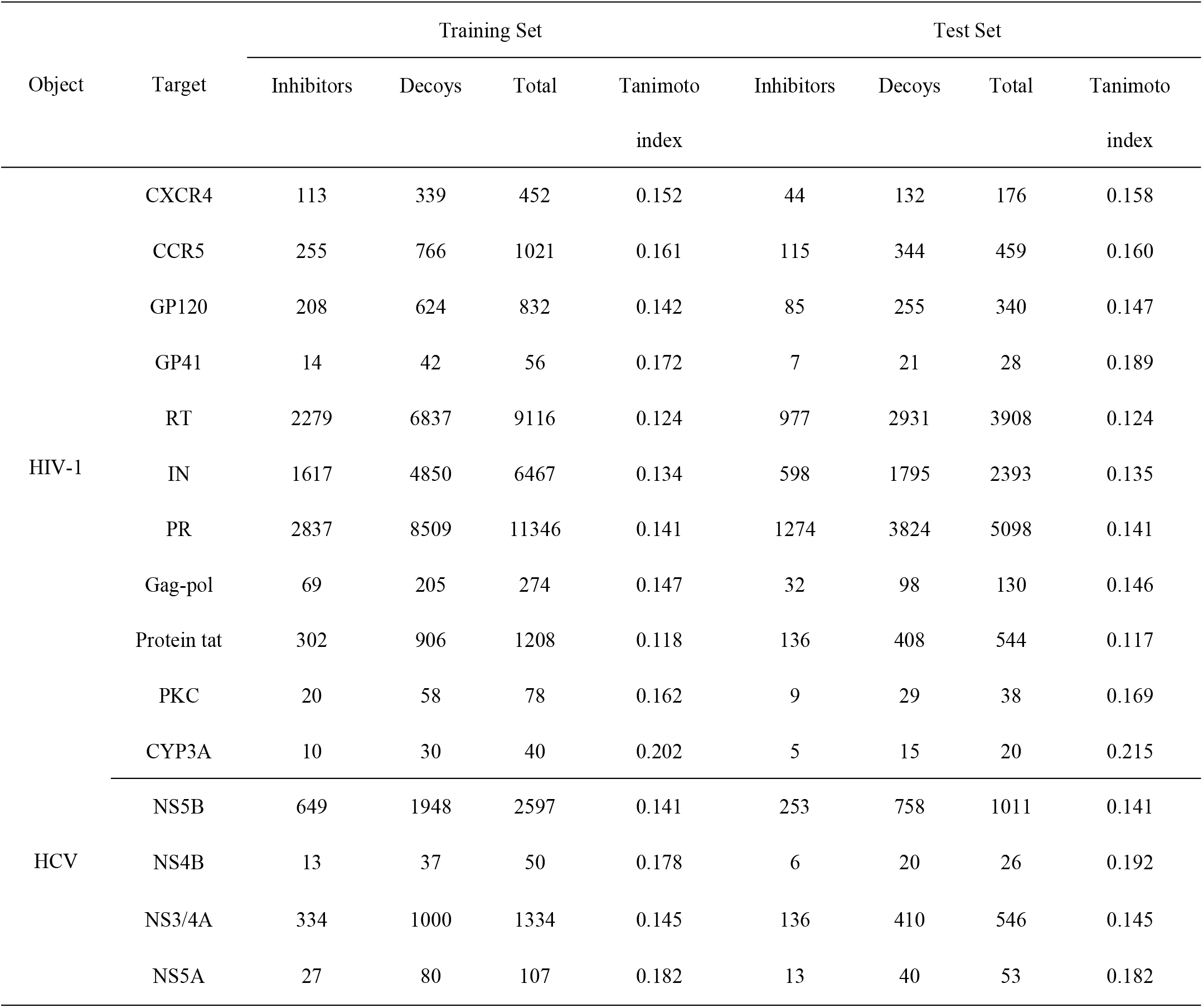
Detailed statistical analysis of the 15 datasets.

**Fig 3.**
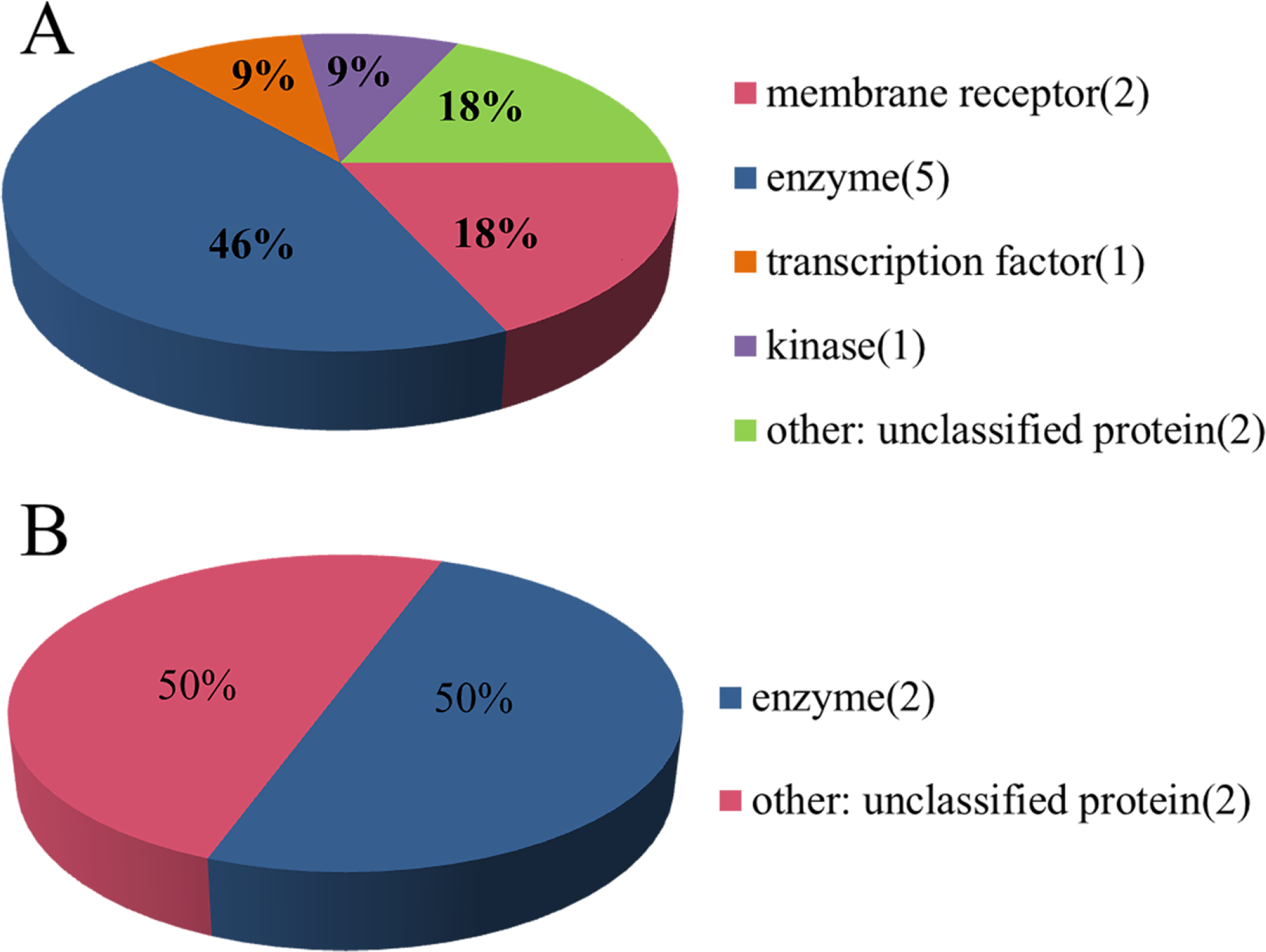
Targets classification of the HIV-1 (A) and HCV (B) data sets.

In addition, according to the protein target classification rules in ChEMBL database, 11 targets related to HIV-1 and 4 targets related to HCV are spatial distributed into five and two categories, respectively. As showed in Fig 3A, the HIV-1 target space (n = 11) consists of five subfamilies, including membrane receptor (n = 2), enzyme (n = 5), transcription factor (n = 1), kinase (n = 1) and unclassified protein (n = 2). Similarly, the HCV target space (n = 4) is divided into two subfamilies, including enzyme (n = 2) and unclassified protein (n = 2) in Fig 3B. Thus, the two data sets for HIV-1 and HCV have diverse target coverage.

### 2.2 Performance evaluation and comparison of models

The construction of all classification models, in this study, was primarily based on two classifiers (NB and SVM) and two types of fingerprints (MACCS and ECFP6). Afterwards, internal 5-fold cross-validation and external test set validation was executed in turn.

The 5-fold cross-validation test results of mt-QSAR models identifying inhibitors and decoys are displayed in Table 2. In the 44 models of HIV-1, 86.36% models (38 out of 44) have AUC value exceeding 0.9, and 61.18% models (30 out of 44) have MCC value exceeding 0.8. In addition, AUC value ranges from 0.83 to 1, with an average of 0.96, and MCC value ranges from 0.52 to 1, with an average of 0.84. Among the 16 models of HCV, all models have AUC value exceeding 0.9, and 81.25% models (13 out of 16) have MCC value exceeding 0.8. Besides, AUC value ranges from 0.92 to 1, with an average of 0.99, and MCC value ranges from 0.52 to 1, with an average of 0.91. Thus, these results suggest that the mt-QSAR models, both for HIV-1 and HCV, have remarkable capability in identifying inhibitors at low false positive rates. Meanwhile, the detailed performance for the training set is listed in S1 Table. Similarly, in the models of HIV-1, 75% models (33 out of 44) have SE value exceeding 0.8, with an average of 0.86, and 95.35% models (41 out of 44) have SP value exceeding 0.8, with an average of 0.96. In the models of HCV, 87.5% models (14 out of 16) have SE value exceeding 0.8, with an average of 0.95, and 93.8% models (15 out of 16) have SP value exceeding 0.8, with an average of 0.96.

**Table 2.**
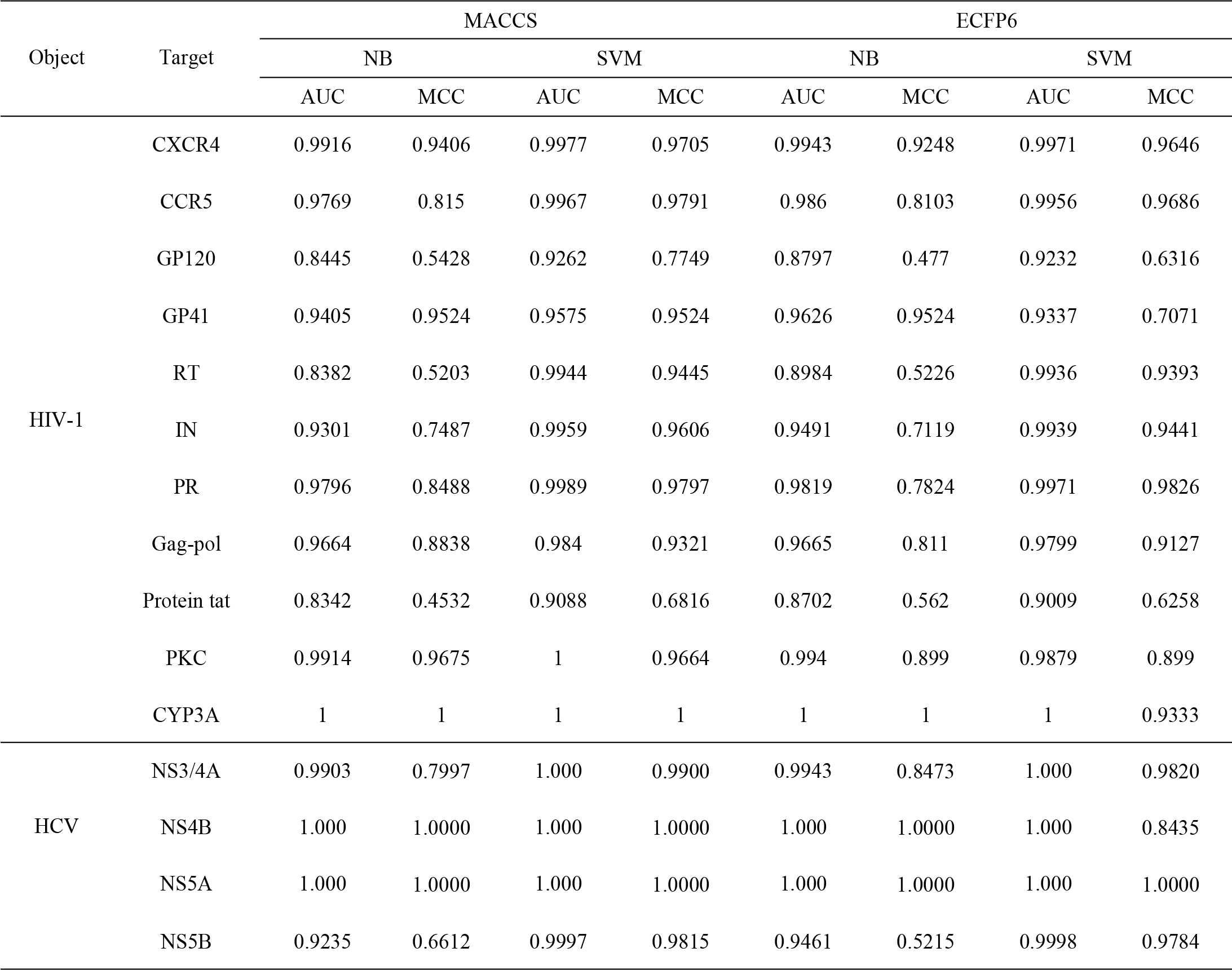
Results of 60 mt-QSAR models’ performance by 5-fold cross-validation.

To assess the generality of the mt-QSAR models on distinguishing inhibitors from decoys, an external test sets were used to perform a more rigorous validation of the training models. The assessment results of test sets are listed in Table 3. In the 44 models of HIV-1, AUC value ranges from 0.83 to 1, with the mean value of 0.97, and MCC value ranges from 0.48 to 1, with the mean value of 0.85. In the 16 models of HCV, AUC value ranges from 0.84 to 1, with the mean value of 0.96, and MCC value ranges from 0.5 to 1, with the mean value of 0.89. The detailed performance of test sets validation is presented in S2 Table. The average value of SE and SP for 11 datasets of HIV-1 is 0.88 and 0.96, respectively. The average value of SE and SP for 4 datasets of HCV is 0.92 and 0.96, respectively. What’s more, prediction accuracy (Q) is an important index for evaluating the predictive ability of the models. Among the models of HIV-1 and HCV, there are 93.18% and 93.75% models with an accuracy higher than 0.8, respectively. According to the results of test sets, the comparable performance of the mt-QSAR models in the training and test sets indicated that the models showed no obviously overfitting. Thus, 60 mt-QSAR models have substantial capability in distinguishing inhibitors from decoys at comparable yield and with low false positive rate.

**Table 3.**
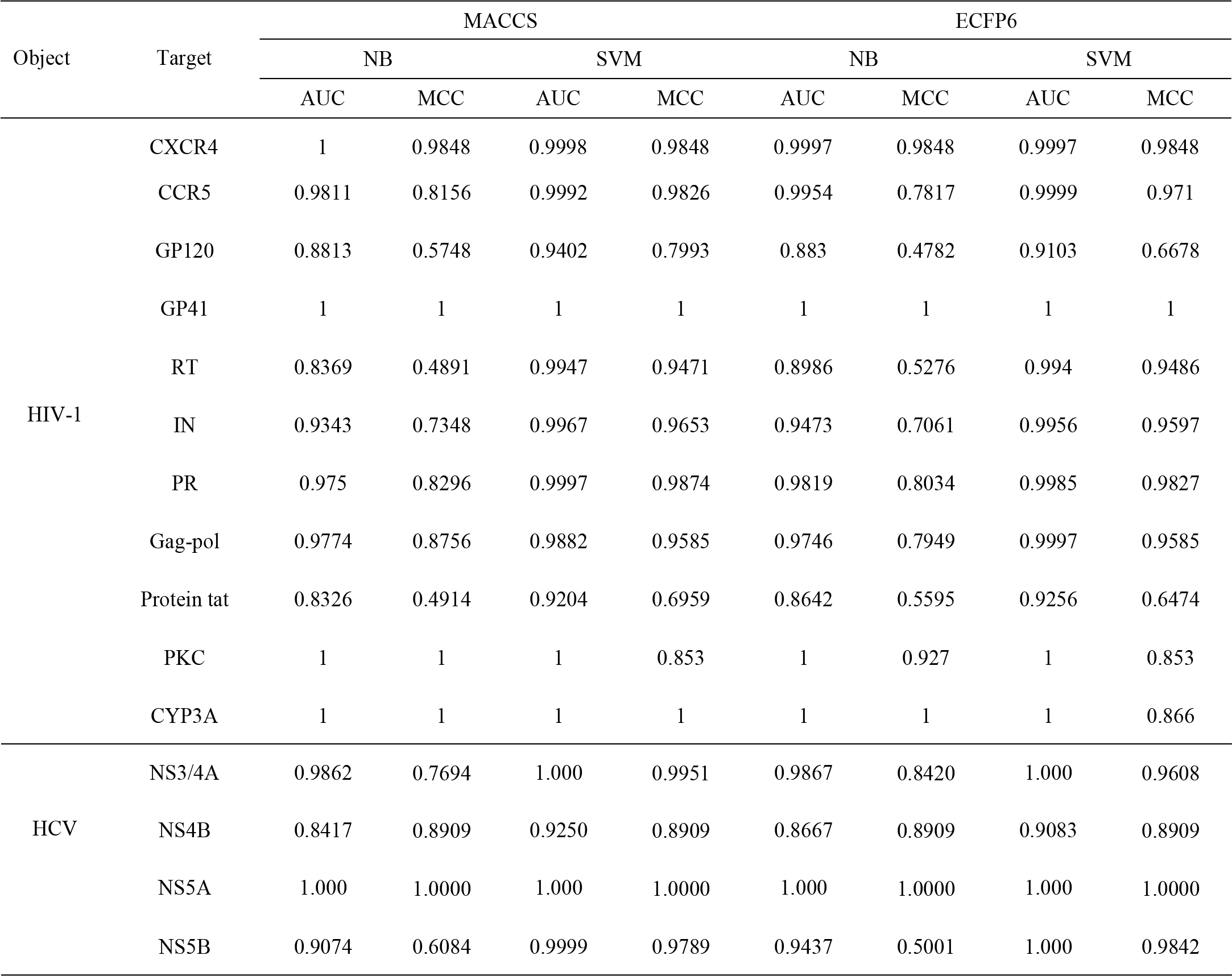
The test results of 60 mt-QSAR models’ performance by test set validation.

To investigate the influence of different types of molecular fingerprints (MACCS and ECFP6) on the performance of naive Bayesian and support vector machine models, MCC values for the test set were compared. As shown in Fig 4, whether for HIV-1 or HCV, MCC values in various mt-QSAR models didn’t differ significantly between MACCS and ECFP6. It should be noted that different fingerprints present different structural representation of a chemical structure, mt-QSAR models based on different fingerprints could improve performance on yield and reducing false positive rate.

**Fig 4.**
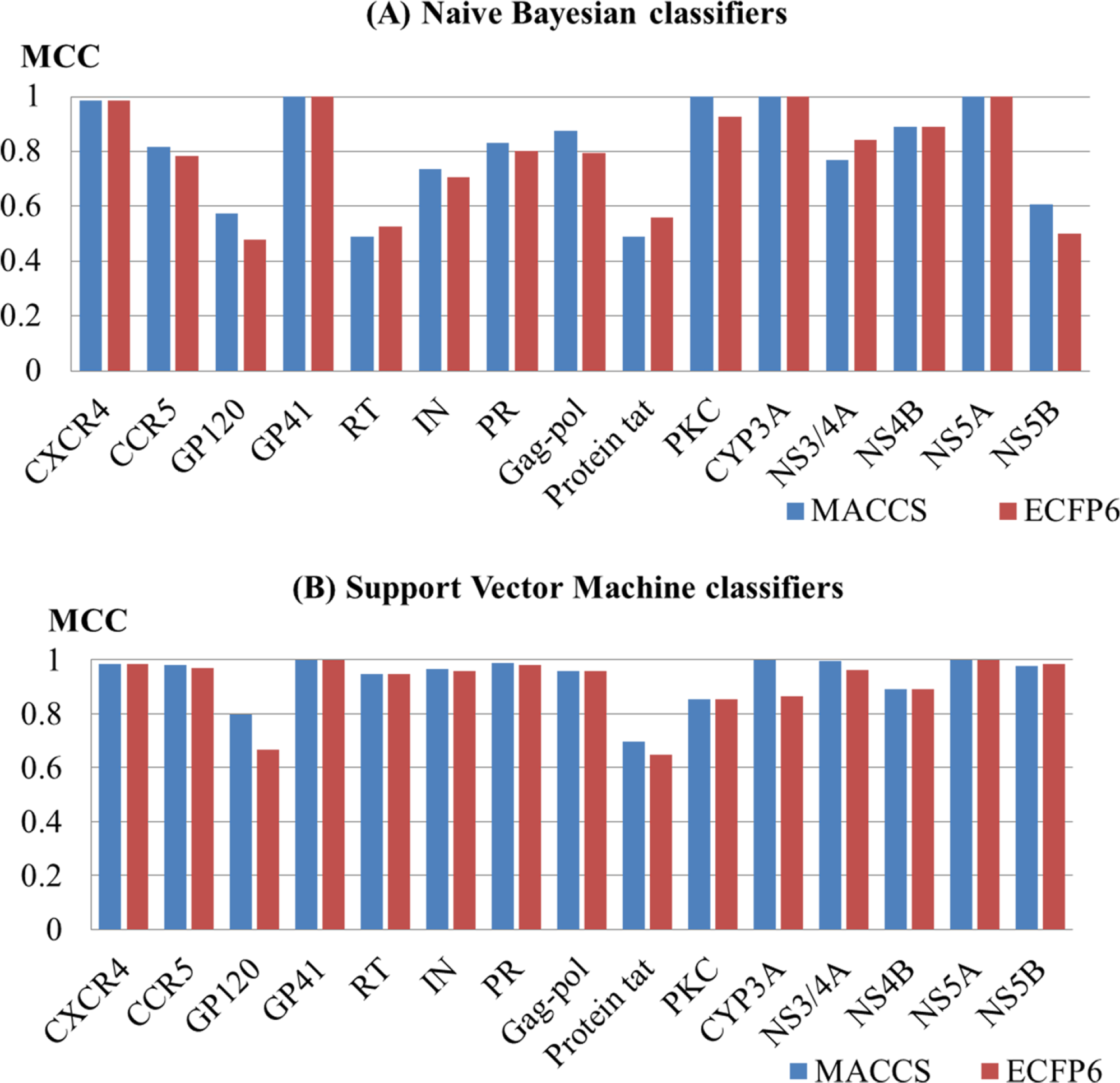
Performance of 60 mt-QSAR models built based on naive Bayesian (A) and support vector machine (B) using different fingerprints by test set validation.

Moreover, the performance of different algorithms (NB and SVM) was also evaluated. With the fingerprint MACCS (Fig 5A), performance of the SVM models are generally superior to that of NB models. In particular, for the models of HIV-1, MCC value from SVM models ranges from 0.70 to 1 with an average of 0.92, better than that from NB models which ranges from 0.49 to 1 with an average of 0.8. For the models of HCV, MCC value from the SVM models ranges from 0.89 to 1 with an average of 0.97, better than that of NB models which ranges from 0.61 to 1 with an average value of 0.82. With the fingerprint ECFP6 (Fig 5B), the models from SVM broadly outperform those from NB. For the models of HIV-1, MCC value from SVM models ranges from 0.65 to 1 with an average of 0.89, better than that from NB which ranges from 0.48 to 1 with an average of 0.78. For the HCV models, MCC value from the SVM ranges from 0.89 to 1 with an average of 0.96, which is dissimilar to NB that ranges from 0.50 to 1 and an average value of 0.81.

**Fig 5.**
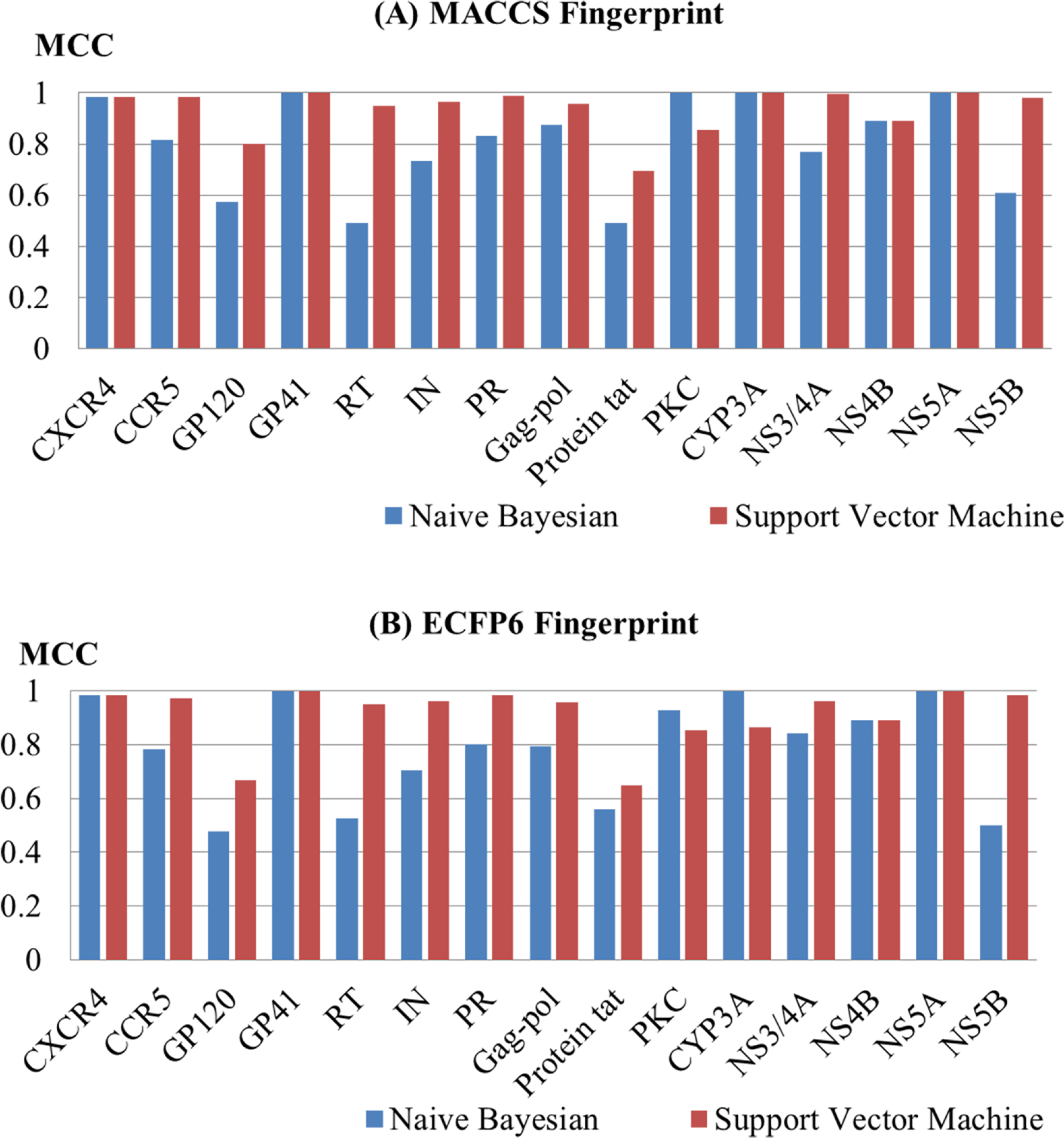
Performance of 60 mt-QSAR models built based on MACCS (A) and ECFP6 (B) using different algorithms by test set validation.

As described above, two fingerprints have no significant difference and the SVM algorithm is slightly better than the NB. However, it is theoretically difficult to determine which fingerprint or algorithm is better because each algorithm or fingerprint has its own advantages and disadvantages. For example, the models built by different fingerprints or algorithms may have opposite predictions for the same molecule. Therefore, combining the results of a single classifier to predict CPIs is indispensable. In this study, four classifiers (NB_MACCS, NB_ECFP6, SVM_MACCS, SVM_ECFP6) were built for each target. One compound was predicted as positive when at least two of four single classifiers predict it to be active, and the corresponding CPIs were defined as potential interactions.

### 2.3 Case 1: Prediction and analysis of polypharmacology for known HIV-1 and HCV drugs

To explore the multiple bioactivities of known drugs, 60 mt-QSAR models based on 15 key targets were used to predict potential targets for 27 approved HIV-1 drugs and 10 approved HCV drugs. If the drug was predicted as active by at least two of the above four classifiers for each target, then the relevant CPI was defined as potential interactions. The detailed prediction results of polypharmacology are presented in S3 Table. The results show that 17 approved HIV-1 drugs and 4 approved HCV drugs were predicted to interact with more than one target, of which 7 approved HIV-1 drugs and 4 approved HCV drugs were predicted to interact with HIV-1 and HCV targets simultaneously. To verify the results of prediction, the predicted targets for each drug were validated using the PubChem BioAssay database[26] and ChEMBL database, and the corresponding binding values are presented in Table 4. Fig 6 shows the histogram of the number of the predicted targets and established targets for 17 HIV-1 approved drugs and 4 HCV approved drugs. For example, Elvitegravir, which was predicted to be active against HIV-1 RT, HIV-1 IN and HCV NS5B, could inhibit HIV-1 RT and IN with IC_50_ value of 91 μM and 28 nM, respectively. Tipranavir, which was predicted to be active against HIV-1 GP120, HIV-1 RT, HIV-1 PR and HCV NS5B, could inhibit HIV-1 PR with an IC_50_ value of 0.03 μM. Telaprevir, which was predicted to be active against HIV-1 PR and HCV NS3/4A, could inhibit HCV NS3/4A with an IC_50_ value of 0.2 μM.

**Table 4.** Known interactions among CPIs predicted by mt-QSAR models for approved drugs.

| Drug name | Predicted targets | Known activity | References |
| --- | --- | --- | --- |
| Enfuvirtide | GP41 | $IC_{50}$ = 90.2 nM | [27] |
| Delavirdine | RT | $IC_{50}$ = 0.17 μM | [28] |
| Didanosine | RT | $IC_{50}$ = 4.6 μM | [29] |
| Lamivudine | RT | $IC_{50}$ = 0.60 μM | [29] |
| Stavudine | RT | $IC_{50}$ = 1.9 μM | [29] |
| Zalcitabine | RT | $IC_{50}$ = 0.13 μM | [29] |
| Zidovudine | RT | IC<sub>50</sub> = 0.13 μM | [29] |
| Dolutegravir | RT | Inactive | [30] |
| Dolutegravir | IN | IC<sub>50</sub> = 2.7 nM | [31] |
| Elvitegravir | RT | IC<sub>50</sub> = 91 μM | [32] |
| Elvitegravir | IN | IC<sub>50</sub> = 28 nM | [32] |
| Amprenavir | PR | IC<sub>50</sub> = 0.15 μM | [33] |
| Atazanavir | RT | IC<sub>50</sub> = 26 nM | [34] |
| Atazanavir | PR | IC<sub>50</sub> = 3.5 nM | [35] |
| Darunavir | PR | IC<sub>50</sub> = 3.0 nM | [35] |
| Indinavir | RT | Inconclusive | [36] |
| Indinavir | PR | IC<sub>50</sub> < 0.01 μM | [36] |
| Nelfinavir | PR | IC<sub>50</sub> = 0.53 nM | [37] |
| Ritonavir | PR | IC<sub>50</sub> = 0.6 nM | [38] |
| Ritonavir | CYP3A | IC<sub>50</sub> = 0.11 μM | [38] |
| Tipranavir | PR | IC<sub>50</sub> = 0.03 μM | [39] |
| Cobicistat | PR | IC<sub>50</sub> = 0.15 μM | [38] |
| Cobicistat | CYP3A | IC<sub>50</sub> > 30 μM | [38] |
| Telaprevir | NS3/4A | IC<sub>50</sub> = 0.2 μM | [40] |
| Simeprevir | NS3/4A | IC<sub>50</sub> = 10 nM | [41] |
| Grazoprevir | NS3/4A | IC<sub>50</sub> = 0.07 nM | [42] |
| Dasabuvir | NS5B | IC<sub>50</sub> = 0.6 μM | [43] |

**Fig 6.**
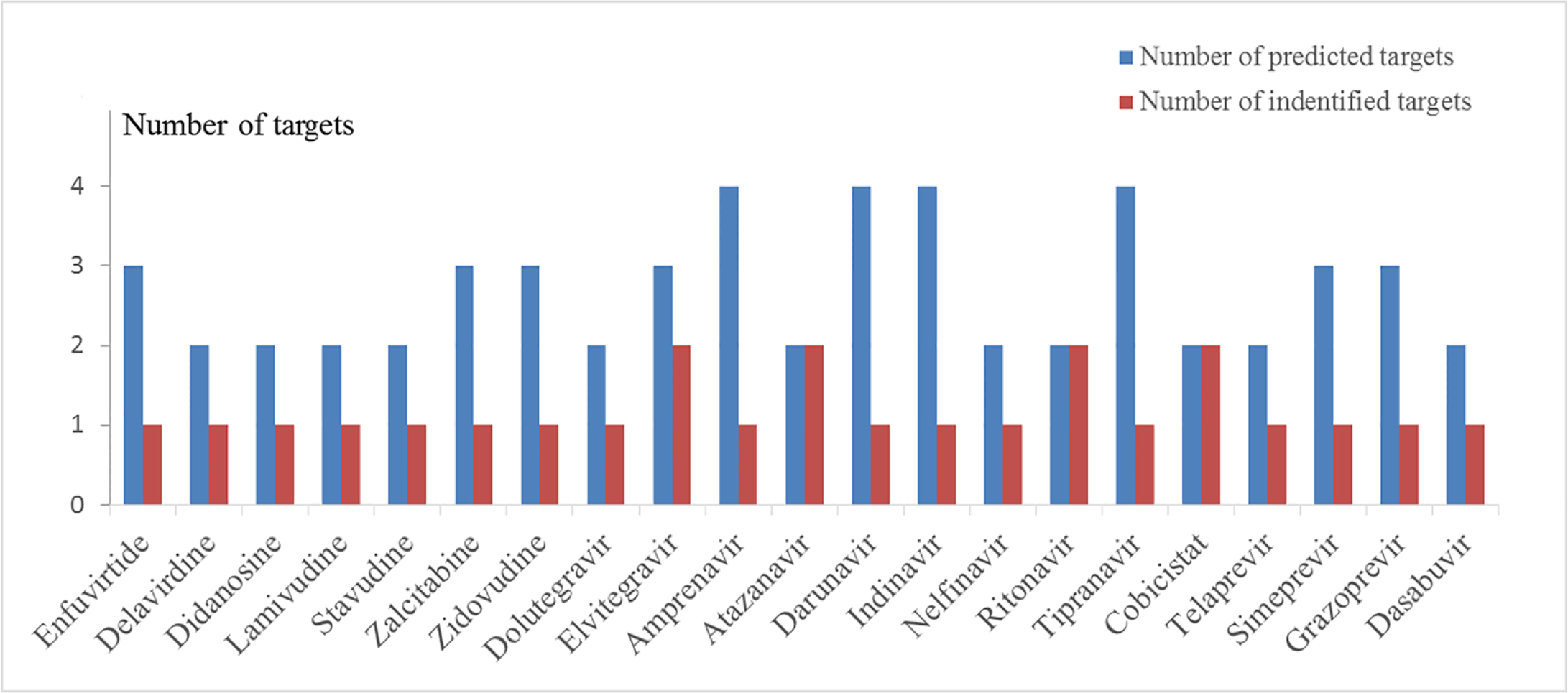
Number of predicted targets and that of known targets for 17 HIV-1 approved drugs and 4 HCV approved drugs.

In addition, 56 CPIs were obtained as shown in S4 Table. Among them, 25 out of 56 predicted CPIs are consistent with experimental results; the remaining 2 CPIs were predicted to be active but were experimentally validated as “Inconclusive” and “Inactive”, respectively[27,28,37–43,29–36]. The success rate of 44.6% (25/56) and the failure rate of 1.8% (1/56) illustrate the reliability of mt-QSAR method. A polypharmacological interaction network between 21 drugs and predicted targets (56 CPIs) were constructed with Cytoscape 3.6[44] (Fig 7). The Fig 7 shows that each drug is predicted to have potential activity on two or three targets, which is in line with concept “one target can hit multiple targets”. Then, predicted CPIs network identifies that many potential drug-target interactions have yet to be explored and unverified CPIs are worthy of further experimental validation. The predicted targets might be new targets for approved drugs and provide a promising therapy for HIV/HCV co-infection.

**Fig 7.**
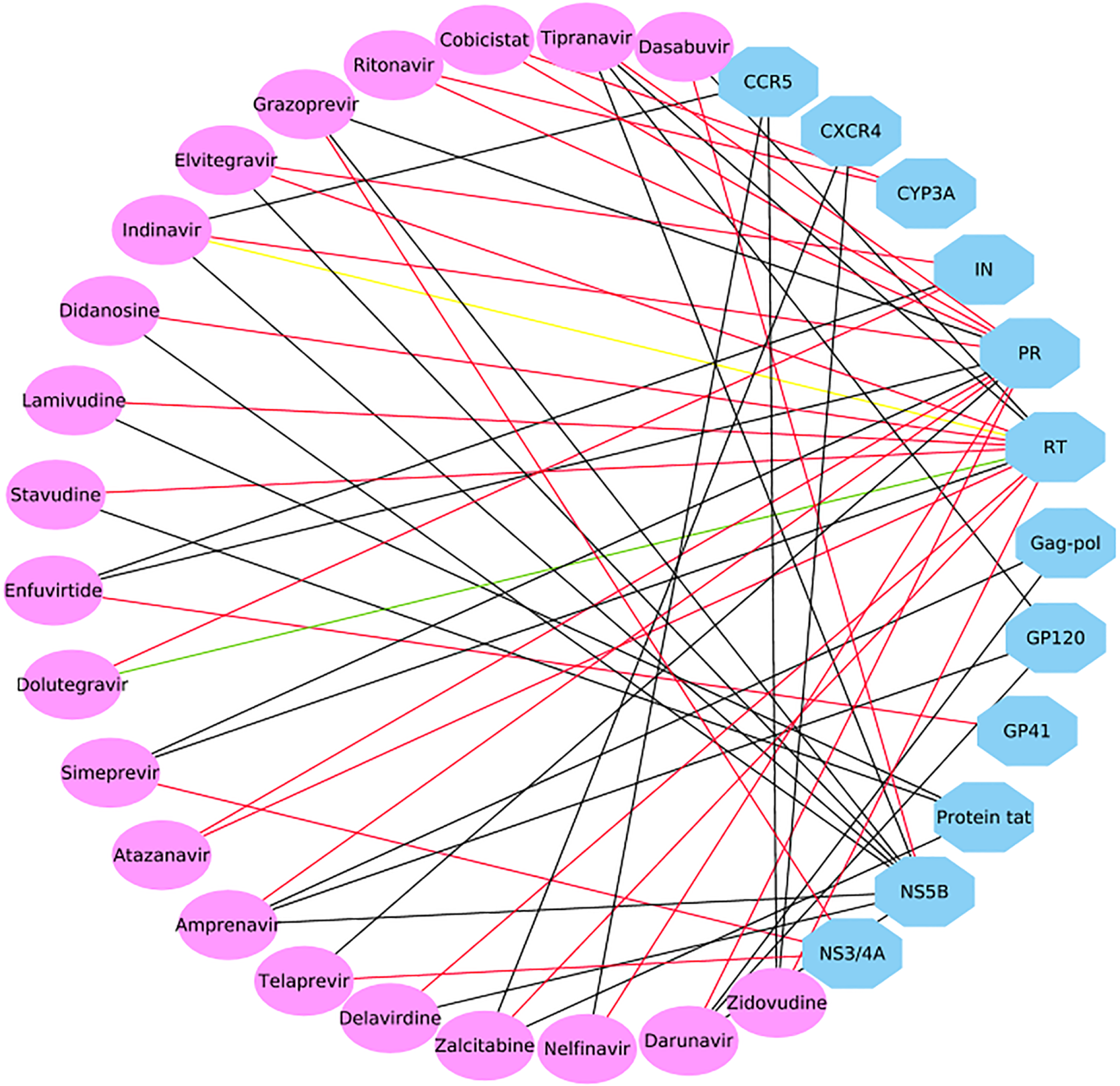
Polypharmacology analysis of 17 HIV-1 approved drugs and 4 HCV approved drugs based on the 60 mt-QSAR models in this study. Ellipses and octagons represent drug nodes and protein nodes, respectively. The red lines indicate that the CPI was experimentally verified as active. The black lines present that the interaction was not proved and worthy of further experimental validation. And the green and yellow lines express that the interaction was experimentally validated as inactive and inconclusive, respectively.

### 2.4 Case 2: Target prediction and analysis for known inhibitors

HIV-1 protease (HIV-1 PR), reverse transcriptase (RT) and integrase (IN) play a pivotal role in the viral replication so that they have become attractive targets to design inhibitors[45]. HIV-1 protease plays a key role in the normal assembly and maturation of virus[46]. HIV-1 reverse transcriptase and integrase catalyze sequentially the synthesis and integration of proviral DNA into the host genome[5]. At the same time, HCV NS5B polymerase is an essential protein involved in the replication of viral positive strand genomic RNA and used as a key target for therapeutic intervention[47,48].

Above all, a set of 9 compounds (Fig 8) with known activities targeting at least one out of four targets (HIV-1 PR, RT, IN and HCV NS5B) and not involved in generating mt-QSAR models were retrieved from ChEMBL database. In order to evaluate whether the new models could correctly differentiate these active compounds, the classifiers built in this study were used to predict chemicals toward HIV-1 PR, RT and IN as well as HCV NS5B. Following the above rules, CPI is referred to a potential interaction when at least two out of four single classifiers predict a compound as positive. The specific predicted results are given in S5 Table, and the comparison of predicted and experimental results is shown in Table 5.

**Table 5.** Predicted and experimental activity of 9 selected compounds towards HIV-1 PR, RT, IN and HCV NS5B.

| CHEMBL ID | PR/Pred<sup>a</sup> | PR/Exp<sup>b</sup> | IN/Pred<sup>c</sup> | IN/Exp<sup>d</sup> | RT/Pred<sup>e</sup> | RT/Exp<sup>f</sup> | NS5B/Pred<sup>g</sup> | NS5B/Exp<sup>h</sup> |
| --- | --- | --- | --- | --- | --- | --- | --- | --- |
| CHEMBL16326 | false | —<sup>i</sup> | true | active | false | — | true | active |
| CHEMBL18927 | false | — | true | active | false | — | true | active |
| CHEMBL449221 | false | — | true | active | false | — | true | active |
| CHEMBL502238 | false | — | true | active | false | — | true | active |
| CHEMBL19332 | false | — | true | active | false | active | true | active |
| CHEMBL210593 | false | — | false | — | true | active | true | active |
| CHEMBL37541 | true | active | false | — | false | — | true | active |
| CHEMBL3612421 | true | active | false | — | false | — | true | — |
| CHEMBL1668670 | false | — | true | active | true | active | true | — |

<sup>a</sup>“PR/Pred” shows the predicted result of a compound against PR. <sup>b</sup>“PR/Exp” shows the experimental result of a compound against PR. <sup>c</sup>“IN/Pred” shows the predicted result of a compound against IN. <sup>d</sup>“IN/Exp” shows the experimental result of a compound against IN. <sup>e</sup>“RT/Pred” shows the predicted result of a compound against RT. <sup>f</sup>“RT/Exp” shows the experimental result of a compound against RT. <sup>g</sup>“NS5B/Pred” shows the predicted result of a compound against NS5B. <sup>h</sup>“NS5B/Exp” shows the experimental result of a compound against NS5B. <sup>i</sup>“—” represents that the predicted result has not been verified by experiments.

**Fig 8.**
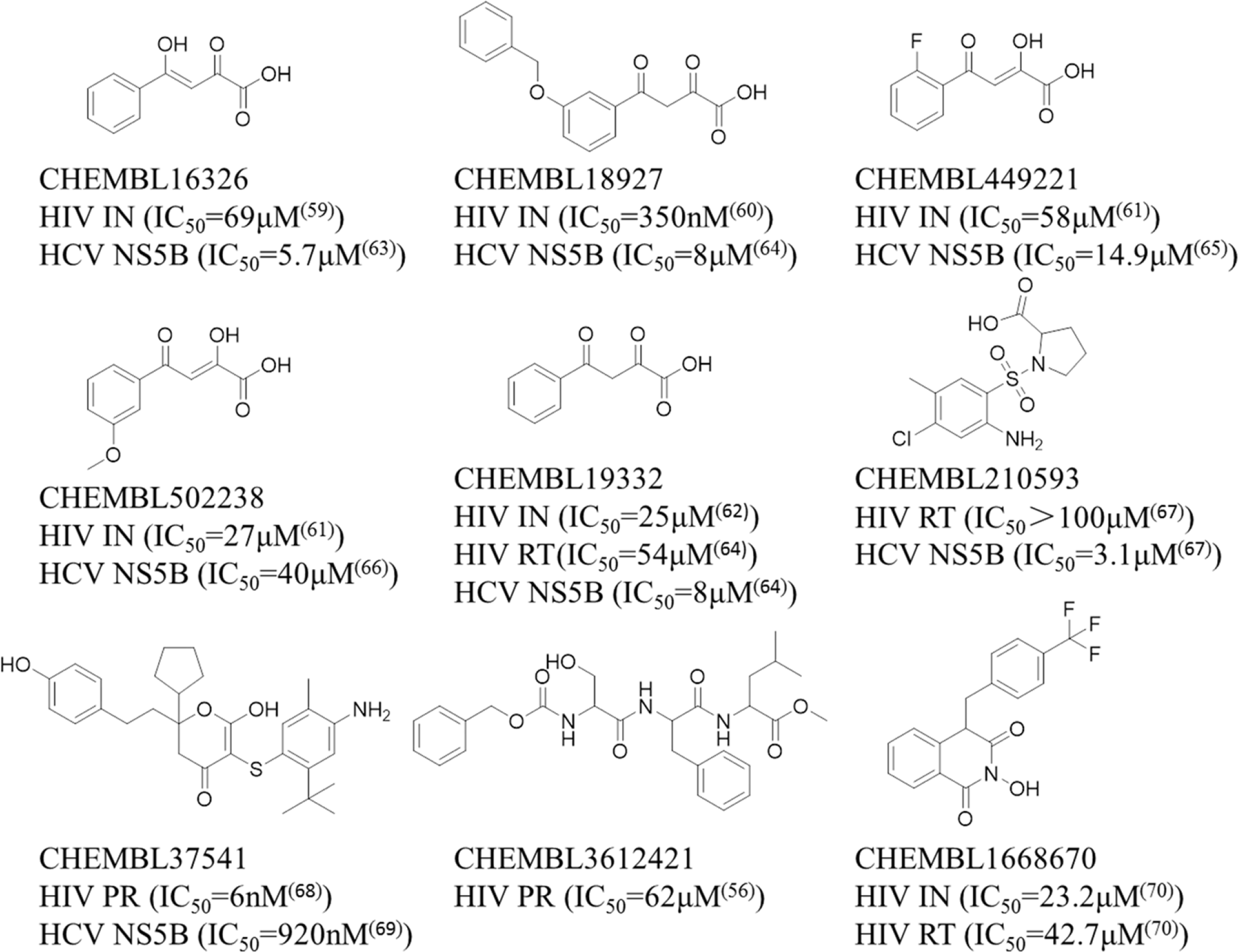
The structure and bioactivity data of 9 selected compounds.

The results in Table 5 show that 5 compounds (CHEMBL16326, CHEMBL18927, CHEMBL449221, CHEMBL502238 and CHEMBL19332) were predicted to be potentially active toward HIV-1 IN and HCV NS5B, and they were also experimentally confirmed to be HIV-1 IN and HCV NS5B inhibitors in the literatures[49–56]. Among them, as a known active HIV-1 RT inhibitor, CHEMBL19332 was shown to be inactive toward HIV-1 RT in the prediction result. In particular, 5 compounds with analogous structures have similar protein-ligand interactions in the active region of HIV-1 IN and HCV NS5B. Moreover, molecular docking provides a more detailed inspection of binding mode for each compound. And the scoring values of the top scoring poses are available in S6 Table. As shown in Fig 9, the hydrophobic groups of those five compounds, such as benzene ring of CHEMBL16326 and monofluorobenzene ring of CHEMBL449221, are placed in the hydrophobic pockets formed by Leu102, Ala128, Trp132, Thr174 and Met178 of HIV-1 IN (see Fig 9A), and formed by Val52, Leu159 and Ile160 of HCV NS5B (see Fig 9B). The diketo acids moiety of those five compounds form hydrogen bonds with Glu170 and His171 in the active site of HIV-1 IN (see Fig 9A), and form hydrogen bonds with Phe224 and Asp225 and metal chelation with Mn^2+^ ions in the catalytic activity site of HCV NS5B (see Fig 9B). Thus, HIV-1 IN and HCV NS5B can be affected by diketo acids inhibitors, which disturbed enaymatic function during the process of virus replication.

**Fig 9.**
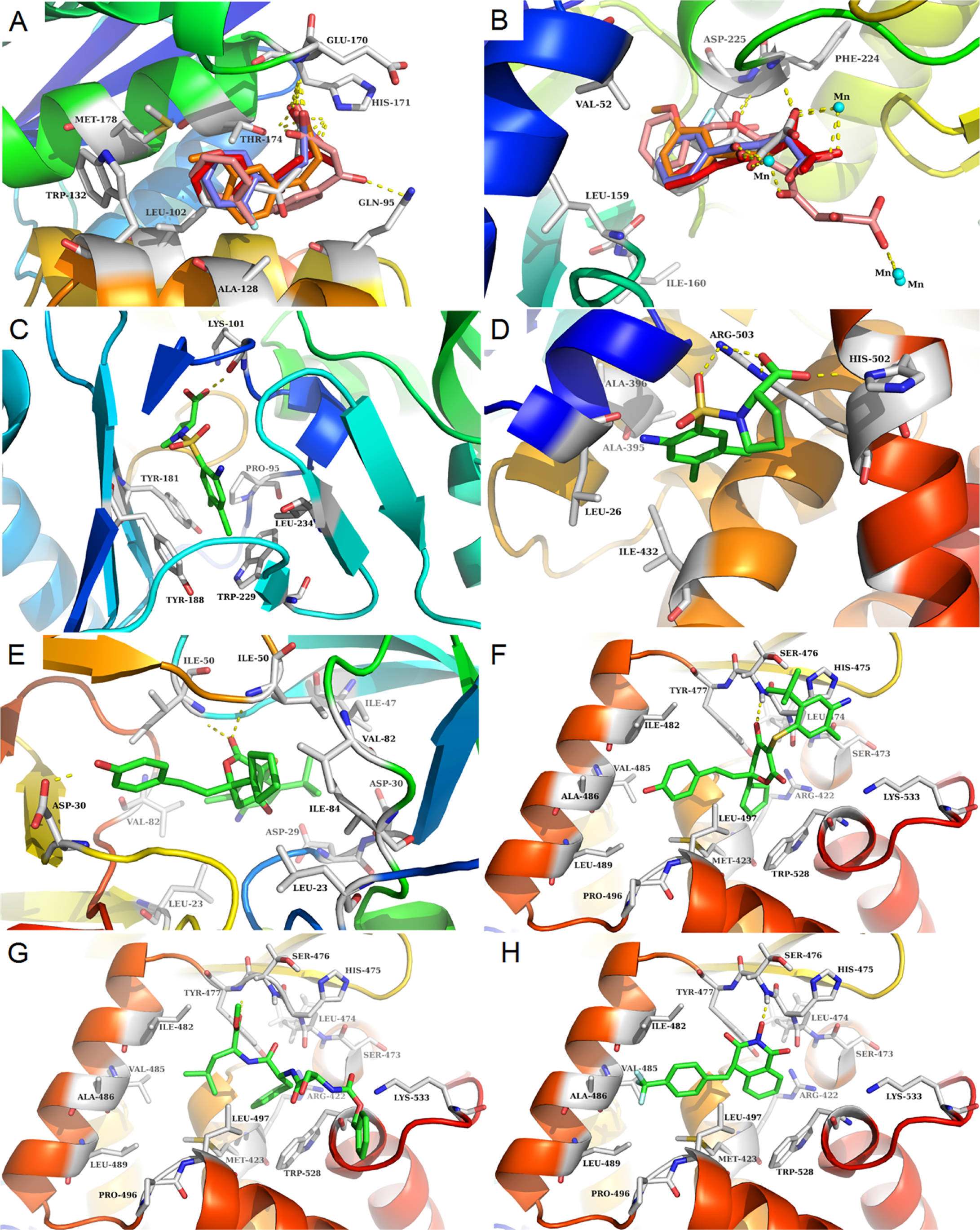
The binding modes of 9 compounds in the respective binding site. A and B present the binding modes of 5 compounds with HIV-1 IN (PDB: 4NYF) and HCV NS5B (PDB: 1GX6), respectively, including CHEMBL16326 (red), CHEMBL18927 (pink), CHEMBL449221 (purple), CHEMBL502238 (orange) and CHEMBL19332 (grey). C and D present the binding modes of the compound (CHEMBL210593) with HIV-1 RT (PDB: 1TL3) and HCV NS5B (PDB: 1OS5). E and F present the binding modes of the compound (CHEMBL37541) with HIV-1 PR (PDB: 4MC1) and HCV NS5B (PDB: 1OS5). G and H present the binding modes of two compounds (CHEMBL3612421 and CHEMBL1668670) with HCV NS5B (PDB 1OS5), respectively.

CHEMBL210593 was predicted as HIV-1 RT and HCV NS5B inhibitors, and it was precisely HIV-1 RT and HCV NS5B inhibitors according to the previous report[57]. Fig 9 shows that 3-chloro-p-methylaniline of CHEMBL210593 occupies the hydrophobic pockets formed by Tyr181, Tyr188 and Trp229 of HIV-1 RT (see Fig 9C), and formed by Leu26, Ala395, Ala396 and Ile432 of HCV NS5B (see Fig 9D). The carboxyl group of CHEMBL210593 forms hydrogen bonds with the main chain carbonyl oxygen of Lys101 of HIV-1 RT (see Fig 9C), and the side chain of Arg503 and His502 of HCV NS5B (see Fig 9D). Therefore, proline sulfonamides inhibitors can affect HIV-1 RT and HCV NS5B and influence enaymatic function.

CHEMBL37541 and CHEMBL3612421 were also predicted to be active against HIV-1 PR and HCV NS5B. CHEMBL3612421 was known HIV-1 PR inhibitor and CHEMBL37541 was known HIV-1 PR and HCV NS5B inhibitor by experimentally validated[46,58,59]. As for CHEMBL37541, the dihydropyrone moiety makes direct hydrogen bonds interaction with Ile50 of HIV-1 PR (see Fig 9E) and Ser476 of HCV NS5B (see Fig 9F). The 4-ethylphenol and cyclopentane groups fit into hydrophobic pockets consisting of the residues Leu23, Leu23’, Val82, Val82’, Ile84 and Ile84’ of HIV-1 PR (see Fig 9E), and the residues Met423, Ile482, Val485, Ala486, Leu489, Leu497 and Trp528 of HCV NS5B (see Fig 9F). New predicted CPI pair involving CHEMBL3612421 and HCV NS5B is presented in Fig 9G. CHEMBL3612421 forms a hydrogen bond with Tyr477 in thumb II region of HCV NS5B and its benzene ring and dimethyl group are located in hydrophobic region consisting of the residues Met423, Ile482, Val485, Ala486, Leu489, Leu497 and Trp528. Similarly, dihydropyrones inhibitors can affect HIV-1 PR and HCV NS5B and influence enaymatic function. CHEMBL1668670 was also predicted as HIV-1 RT, IN and HCV NS5B inhibitor, and it only have bioactivity data against HIV-1 RT and IN[60]. New predicted CPI pair involving CHEMBL1668670 and NS5B was presented in Fig 9H. CHEMBL1668670 forms a hydrogen bonding with Ser476 in thumb II region of HCV NS5B and its trifluorobenzene ring is directed to the hydrophobic region consisting of the residues Met423, Ile482, Val485, Ala486, Leu489 and Leu497.

Thus, 7 out of 9 compounds, which are known to act on HIV/HCV co-infection targets, were successfully predicted to act on two or three targets. New predictions involving CHEMBL3612421 and CHEMBL1668670 targeting NS5B have yet to be further validated by experimentally. The result is given that design of specific inhibitors targeting HIV-1 targets and HCV targets simultaneously is now possible. In addition, one individual compound has co-inhibitor ability for both virus targets if it is able to satisfy the requirements of binding the HIV/HCV target active site in the molecular shape, size and physical chemical properties. In short, mt-QSAR models for HIV/HCV co-infection are expected to contribute to design and identify different types of co-inhibitors with excellent affinities that selectively and simultaneously target multiple HIV-1 and HCV targets.

In summary, 19 CPI pairs were predicted to be positive, and of which 17 CPI pairs were experimentally confirmed to be active. Therefore, prediction success rate of 89.5% (17/19) indicates the reliability of the mt-QSAR models. It is worth mentioning that remaining 2 CPI pairs related to 2 compounds deserve further investigation by experiments, and the predicted targets might be the new targets for the known inhibitors. Moreover, one CPI pair shows false negative in the prediction result.

The prediction results demonstrate that machine learning models have high specific and low false negative rate in this study. Here, two possible reasons for causing false negative are as follows. Firstly, molecular fingerprint is unable to sufficiently represent molecular structural information. Secondly, the inherent limitations of machine learning and the composition of training set have a significant impact on the accuracy of the models.

## 3. Discussion

In the past, the “one gene, one drug, one disease” was the main paradigm of drug discovery. However, with the introduction of the concept of polypharmacology, multitarget therapy using a single drug that simultaneously inhibits two or more targets may provide a new perspective. Here, the mt-QSAR method was applied to identify the compounds acting on multiple targets for HIV/HCV co-infection.

In this study, based on the mt-QSAR method, 44 binary classifiers of 11 targets related to HIV-1 and 16 binary classifiers of 4 targets related to HCV were established to predict the CPIs. Besides, the prediction reliability of models was confirmed by 5-fold cross-validation and test set validation.

To illustrate the application of the models, two different cases were performed to systematically predict the multiple bioactivities for 27 approved HIV-1 drugs and 10 approved HCV drugs (case 1) and 9 compounds that were known to be active toward at least one target of HIV-1 and HCV (case 2). For case 1, 21 approved drugs were predicted to be active against more than one biological target. And the predictions were confirmed by reported bioactivity data, the success rate was 44.6% and the failure rate was 1.8%. For case 2, 9 active compounds toward at least one target (HIV-1 PR, RT, IN and HCV NS5B) were predicted to be multitarget inhibitors by mt-QSAR models with a success rate of 89.5%. In short, the prediction results of above two cases indicate that mt-QSAR models have a significant performance on prediction of multitarget inhibitors against HIV/HCV co-infection.

As mentioned above, mt-QSAR method achieved satisfactory prediction accuracy, and machine learning models were applicable for potential targets identification in this study. However, similarity searching and molecular docking will be better choices for those targets with a small number of active molecules, such as GP41, PKC, CYP3A, NS4B and NS5A whose active molecules are less than 30. Besides, machine learning models and similarity searching neglect the interaction between receptor and ligands compared with molecular docking. Therefore, it is necessary to choose a reasonable single-method or combining of multi-methods according to specific situation.

In summary, computational methods utilized in this study could effectively detect multitarget co-inhibitors for suppression of HIV/HCV co-infection, and have potential application in identifying new targets, virtually screening of multitarget drugs and drug repurposing in treatment of other diseases. In addition, we hope that our research will be of help to the clinic treatment of HIV/HCV co-infection.

## 4. Materials and methods

### 4.1 Dataset preparation

Therapeutic Target Database (TTD)[61] and ChEMBL Database (version 23, https://www.ebi.ac.uk/chembl/) were used to collect the clinical targets of anti-HIV and anti-HCV therapy. We used keywords with ‘HIV-1’, ‘Human Immunodeficiency Virus type-1’, ‘HCV’ and ‘Hepatitis C Virus’ as the retrieval conditions, and the search results were further filtered to keep the targets with drug candidates that at least entered phase I clinical trials.

In this study, the known small molecule inhibitors of the HIV-1 or HCV targets were downloaded from the ChEMBL Database, respectively. Data sets were processed according to the following requirements: (1) the compound, with IC_50_, EC_50_ or Ki ≤ 20 μM, was retained as positive (expressed as ‘+1’); (2) salts were translated into the homologous acid or base, water molecules in the hydrate were eliminated; (3) the duplicate molecules were deleted. On the other hand, DecoyFinder software[62,63] was applied to prepare a set of decoy compounds (3 decoys for each known inhibitor)[64] from ZINC database (version 12, http://zinc.docking.org/), and decoy compounds were used as negative (expressed as ‘-1’) for each target. Particularly, decoy compounds were selected based on the following constraints: (i) the MW is within 25 Da of positive compounds; (ii) the number of RB, HBDs, HBAs are ±1, ±1 and ±2 of positive compounds, respectively; (iii) the LogP is within 1 of positive compounds; (iv) the Tanimoto coefficients of a potential decoy vs each active compound as well as a potential decoy vs previously selected decoys are the default of 0.75 and 0.9, respectively[62]. Finally, the data set including inhibitors and decoys for each target was divided into training set and test set. The data processing procedures were performed by KNIME (version 3.4.2, https://www.knime.com), an open-source data analysis and cheminformatic software.

### 4.2 Molecular representation

Each compound in the above data sets was represented using two different types of binary fingerprint descriptors (MACCS and ECFP6) as chemical descriptors calculated in KNIME. MACCS is a common 2D molecular fingerprint descriptor, composed of 166 structural keys. And most of the important chemical features are covered in spite of small length[65]. Additionally, Extended-connectivity fingerprints (ECFPs) are a new kind of topological fingerprints, which were clearly developed to capture molecular features associated with molecular activity[23]. At present, MACCS and ECFPs have been widely used in drug discovery on the basis of the characteristic of calculating rapidly and representing a lot of different molecular features and stereochemical information.

### 4.3 mt-QSAR models building

In this study, all classification models were built by combining two machine learning algorithms (naive Bayesian and support vector machine) and two molecular fingerprints (MACCS and ECFP6). In short, four classifiers (NB_MACCS, NB_ECFP6, SVM_MACCS and SVM_ECFP6) were generated for each target. Therefore, the numbers of mt-QSAR models for 11 HIV-1 targets and 4 HCV targets are 44 and 16, respectively.

#### 4.3.1 Naive Bayesian (NB)

The Bayesian classification is widely used to solve classification problems based on Bayesian theory, such as discriminating between active and inactive compounds by building classifier and so on. And naive Bayesian is one of the simplest and the most common classifier, which adopts the hypothesis of attribution conditional independence[19]. In other words, it is assumed that each feature has an effect on the classification result independently. The details of the Bayesian theorem are given in (equations (1) – (3):

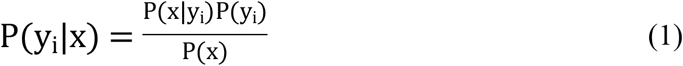

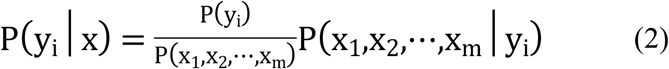

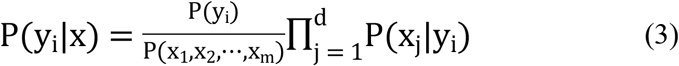

Here, x denotes a set of features such as a molecular fingerprint is the feature set of a molecule; y represents a set of categories that mark the molecular activity category with “+1” or “-1”; x_j_, y_i_ and d are j-th attribute value, i-th category and the amount of attributes, respectively. Besides, P(y_i_) means prior probabilities, which is calculated by analyzing the training set data; P(x_1_,x_2_,…,x_m_) is a constant for the explicit training set.

Because of the assumption of attribution independence, equation 2 is simplified to equation 3, namely the naive Bayesian formula.

#### 4.3.2 Support vector machine (SVM)

Based on the principle of structural risk minimization(SRM)[66], SVM has been developed as a useful classification technique. The main idea is to convert input sample space into a high-dimensional space, in which a hyperplane with the maximal margin is generated to describe non-linear classification boundary. Importantly, the classification is achieved through a kernel function. There are four basic kernels: linear, polynomial, radial basis function (RBF) and sigmoid[67]. In most cases, RBF is preferred[67]. In this study, the RBF kernel function was also used. The penalty parameter C and kernel parameter γ, involved in SVM, were optimized through 5-fold cross-validation.

### 4.4 Performance evaluation of mt-QSAR models

All models were evaluated by the internal 5-fold cross-validation and external test set validation. In 5-fold cross-validation, the training set was divided into 5 subsets equally. Four subsets were used to train the model, and the remaining one was used as an internal validation set.

The major evaluation indexes include true positives (TP), true negatives (TN), false positives (FP), false negatives (FN), sensitivity (SE), specificity (SP), the overall prediction accuracy (Q) and the receiver operating characteristic curve (ROC). Among them, the ROC curves are often used to estimate the merits of a binary classifier, in which the false positive rate (FPR) and true positive rate (TPR) are used as abscissa and ordinate, respectively. Generally, the AUC value ranges from 0 to 1, and the higher the value, the better the model’s classification capability is.

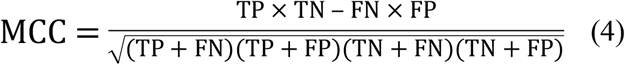

In addition, Matthews correlation coefficient (MCC) is calculated by equation 4. And the value of MCC ranges from −1 to +1.

### 4.5 Molecular docking

The docking algorithm Glide[68] was used for molecular docking and analysis of the binding modes between molecules and target proteins. The crystal structure of HIV-1 IN (PDB: 4NYF)[69], HIV-1 RT (PDB: 1TL3)[70], HIV-1 PR (PDB: 4MC1)[71] and HCV NS5B (PDB: 1OS5 and 1GX6)[72,73] were prepared and used to build the receptor grid. The grid was centered on the ligand of co-crystallized complex or the key residues of proteins, and then the Glide SP protocol was utilized for docking. For the grid generation and docking, all of the parameters are default values.

## Acknowledgments

This study was supported by the National Key R&D Program of China (No. 2017YFC1104400).

